# Red light mitigates ALAN-induced memory impairment in the clownfish *Amphiprion ocellaris*

**DOI:** 10.64898/2026.01.14.699448

**Authors:** Jan IJ van Straaten, Christin Mundell, Fabio Cortesi

## Abstract

Artificial light at night (ALAN) is an increasingly pervasive anthropogenic pollutant in coastal ecosystems. However, its effects on brain function and behaviour in reef fishes remain poorly understood. Here, we investigated the impact of ALAN on learning and memory in the clownfish, *Amphiprion ocellaris*, a diurnal reef fish with well-characterised photoreceptive systems. Cognitive performance was assessed using a colour discrimination task following exposure to ALAN of varying intensities and spectral compositions. ALAN impaired learning and memory after only two consecutive nights of exposure, with effects emerging at intensities as low as 0.5 lux. At higher intensities (5–50 lux), memory was consistently disrupted under white-cool, white-warm, blue, and green LED light, whereas red light had no detectable effect. Together, these findings demonstrate that ALAN rapidly alters cognitive performance in clownfish and that these effects are strongly wavelength-dependent. Our results highlight the vulnerability of diurnal reef-associated fishes to nocturnal light pollution and underscore the need for ecologically informed lighting strategies that use longer-wavelength red-emitting light sources to reduce both the intensity and spectral impact of ALAN in coastal environments.

## Introduction

Artificial light at night (ALAN) is an emerging terrestrial and aquatic anthropogenic pollution worldwide (Tidau et al., 2021). In the marine realm, more than 3.1% of the world’s exclusive economic zones, or 1.9 million km^2^, are already impacted by ALAN, many of which are nearshore marine environments (Smyth et al., 2021). Still, ALAN can also affect offshore environments due to the reflection of light from clouds (skyglow) and shipping (Tidau et al., 2021; Marangoni et al., 2022). The increased use of energy-efficient, short-wavelength ‘white-cool’ LEDs further intensifies the environmental impact, as the emitted shorter blue-to-green wavelengths penetrate deeper into the water and disrupt the circadian rhythms, physiology, and behaviour of marine organisms (Brüning et al., 2018; Li et al., 2024).

While the effect of ALAN on marine invertebrates, such as corals and crustaceans, is well studied (Marangoni et al., 2022), how it affects the ecology and behaviour of marine vertebrates, specifically coral reef fishes, is poorly understood. For example, in the freshwater zebrafish*, Danio rerio*, ALAN alters learning and has task-specific effects on cognitive flexibility (De Russi et al., 2024; Lucon-Xiccato et al., 2023). Similarly, reef fishes rely on cognitive functions such as memory and learning to support critical behaviours, including social interactions, foraging, and predator avoidance (Kieffer & Colgan, 1992). Consequently, ALAN-induced disruptions to cognitive processes may impair these essential behaviours, ultimately affecting reproduction and survival. This is supported by a recent study, which found that ALAN impaired learning in the cleaner wrasse, *Labroides dimidiatus* (Sowersby et al., 2025). However, which aspects of ALAN (i.e., intensity, spectrum, and duration) caused the impairment were not investigated. Understanding the specifics by which ALAN affects brain function and behaviour may have far-reaching implications, e.g., for conservation-oriented strategies to implement ecologically safer light intensities and spectra in coastal environments (Ferretti et al., 2024). Central to ALAN’s perception in the first place are the various light-sensitive organs coral reef fishes possess.

As in other vertebrates, light in reef fishes is captured by photoreceptors, which transmit an electrical signal that initiates the phototransduction cascade, allowing downstream processing (Lamb et al., 2016). Retinal photoreceptors (rods, cones and photoreceptive ganglion cells) support visually guided behaviours such as foraging, predator avoidance, and colour discrimination, as well as entraining circadian rhythms in the eye (Marshall et al., 2019). Extra-retinal photoreceptors, on the other hand, particularly those in the pineal gland, entrain circadian rhythms outside of the eye (Ben-Moshe Livne et al., 2016; Ben-Moshe et al., 2014).

Light-mediated circadian rhythms are well-characterised in zebrafish, where they regulate processes such as melatonin secretion, opsin gene expression, retinal activity, and sleep-wake cycles (Li, 2019; Ben-Moshe Livne et al., 2016). However, their function in reef fishes remains poorly understood. Recent studies on clownfishes (subfamily Amphiprioninae) have described the diversity and distribution of photoreceptors in the retina and pineal gland, highlighting these species as valuable models for examining light sensing and circadian regulation in a reef-fish context (Cortesi et al., 2022; Mitchell et al., 2021). Moreover, clownfishes are emerging model organisms for marine science that can be readily raised in captivity (Laudet & Ravasi, 2023). They are amenable to training and learning (Powell et al., 2021) and are capable of performing complex visual and cognitive tasks (Mitchell et al., 2024), making them ideal for studying the effects of ALAN on cognitive performance.

Here, we examine the effects of ALAN on memory and learning in the false percula clownfish, *Amphiprion ocellaris*. Cognitive performance was assessed using a colour discrimination task (as per Triki et al., 2023), following the exposure over two consecutive nights to different intensities of white-cool LED ALAN (0.5, 5, and 50 lux) and to different colours of ALAN (all at 50 lux). These treatments were selected to compare modern (white-cool LED) versus older-style (white-warm LED) ALAN, and to test a range of wavelengths (blue, red, and green) to identify the spectrum-dependent effects on the brain. We found intensity-dependent effects of ALAN on memory and learning. In addition, while all other tested spectral compositions of ALAN negatively impacted the fish’s cognitive performance, red light had no effect.

## Material & Methods

### Animal husbandry

Darwin morphs (black) false percula clownfish, *Amphiprion ocellaris*, were captive-bred as part of the Marine Sensory Ecology Group’s breeding colony at the Institute of Molecular Biosciences, University of Queensland (offspring from breeding pairs P20 and P6). Approximately 1-year-old individuals were housed in pairs, with the larger fish as the female and the smaller as the male, allowing us to distinguish individuals in each tank by size. Fish were kept under laboratory conditions (broad white-warm LED Lights, 12L:12D, artificial seawater at 1.025 ppm salinity, 27-28 °C) in 80 L tanks and provided with a small terracotta pot as shelter. Before training, fish were acclimated to their experimental aquarium for two weeks. Sixteen fish (eight females and eight males) were used in the intensity experiment, and 16 different fish (eight females and eight males) were used in the experiment testing the different spectral compositions of ALAN. In each experiment, four pairs served as controls and four as treatments; no pair was used in both conditions. Animal care and euthanasia followed procedures approved by the University of Queensland’s Animal Ethics Committee (2022/AE000569).

### Experimental setup and paired-choice colour discrimination test

The experimental setup and paired-choice colour discrimination paradigms were conducted as described in detail by Triki et al. (2023). Briefly, experimental aquaria were partitioned into separate housing and testing compartments using opaque plexiglass (Figure 1). Using this physical and visual separation between a pair of fish prevented social learning during trials (Figure 1). Prior to each test trial, individual fish were confined to a smaller housing compartment within the test compartment using a pair of acrylic barriers—one transparent and one opaque (dimensions: 24 × 22 cm). The removal of the opaque barrier, followed by the transparent barrier, enabled the fish to visually inspect the test compartment before engaging with the task.

**Figure 1:**
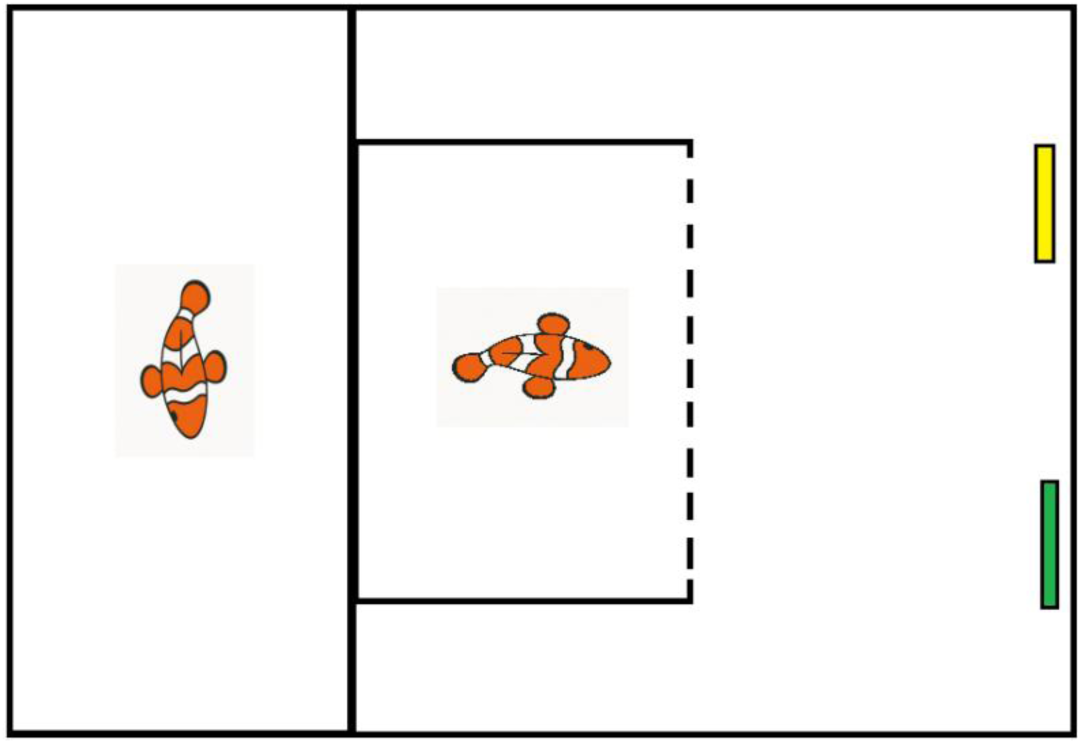
Overview of the experimental setup. Fish were divided into the holding compartment (left fish) and the housing compartment within the test compartment (right fish), with the compartment and the removal barrier made from opaque acrylic. For the two-choice experiment, a yellow (reward) and green cue were placed at the far end of the test compartment. The test fish was kept in the test compartment, with the dotted line representing two acrylic barriers, one transparent and one opaque. The opaque barrier was initially lifted to allow the fish to become accustomed to the test arena before the transparent barrier was lifted.

Before training, fish were habituated to isolation in the test compartment, then to feeding on shrimp paste (approx. 0.5 cm in diameter) smeared onto grey plastic plates (5cm x 5cm 0.5cm), and subsequently to feeding from the designated colour cue (coloured plastic plates with the same dimensions as the grey ones). During training and trials, fish were presented with a yellow and green cue, which was taped to the grey plastic support (Figure 1). We used yellow as the rewarding cue, with shrimp paste stuck to the back of the support, invisible to the fish, and green as the control. Following cue placement, the opaque barrier was removed to allow visual inspection of the test compartment. After two seconds, the acrylic sheet was lifted, permitting the fish to engage in the task (Fig. 1). To claim their reward, fish had to swim to the back of the yellow cue. A fish approaching the yellow cue and claiming the reward was scored a “success”, while a fish that first explored the green cue was scored a “failure”, which resulted in cues being removed. Fish were trained for 3 weeks to learn the task, and testing began when both the treatment and control groups exceeded 85% success across 10 trials. These learning criteria correspond to a probability of correct responses exceeding the chance level of 50% (*p* < 0.05, binomial test). During the testing phase, each fish underwent 10 trials per day over 3 consecutive days (2 nights of ALAN exposure). After three days of testing, the fish were left to recover under normal dark-light conditions for two days. Then, the fish underwent two days of reinforcement training in the colour discrimination task to restore the 85% threshold before the next testing phase began.

### Light treatment

For the intensity experiment, ALAN was generated using broad-spectrum white-blue LED lights (VIPARSPECTRA, Model V165) at three intensities: 0.5, 5 and 50 lux as measured with a Lutron LX-105 light meter (https://lutron.com.tw). For 0.5 lux, the LED light was set to 1/100; for 5 lux, to 4/100; and for the 50 lux treatment, to 15/100. The light intensity was measured at the water surface in the centre of the tank, and fish were exposed to ALAN from 7 p.m. to 7 a.m. Based on the effects of these three intensities (Fig. 2), 50 lux was chosen as the standard intensity for comparing the impact of different spectral compositions of ALAN on memory and learning in *A. ocellaris*.

**Figure 2:**
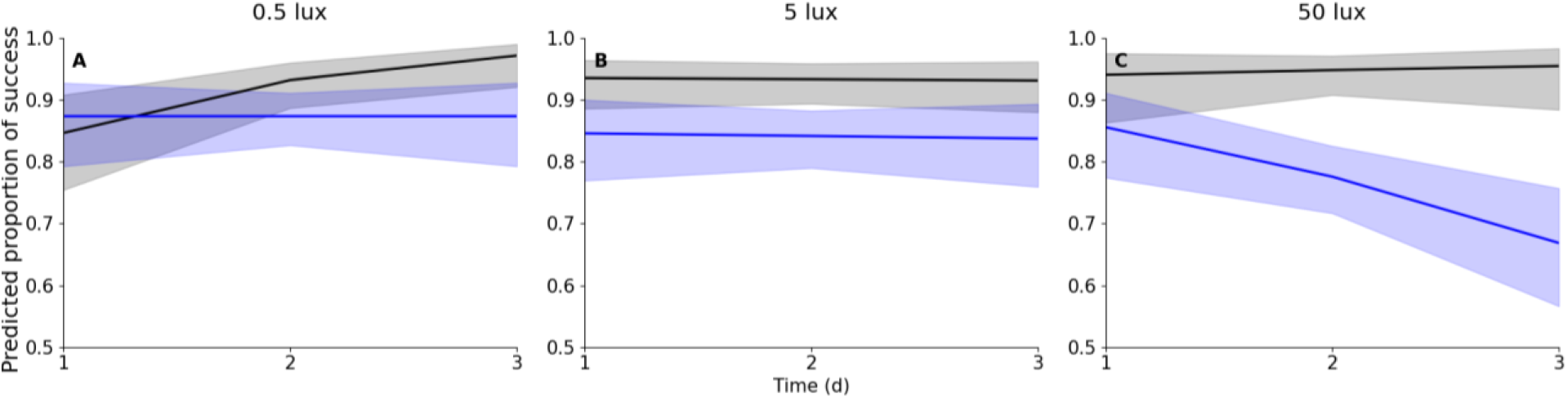
The impact of different intensities of white-cool LED light at night on *Amphiprion ocellaris* memory and learning. Each panel shows the model-predicted mean (solid line) and 95% confidence interval (shaded area) for fish exposed to ALAN over two consecutive nights at 0.5 (A), 5 (B), or 50 lux (C), compared to the dark control group (black) (n = 8 fish per treatment). The y-axis starts at 0.5, representing random chance. See Figs. 3 and S1 for the spectral composition of the white-cool LED at 50 lux.

To test the effects of ALAN with different spectral compositions, we used broad-spectrum LED lights (VIPARSPECTRA, Model V165) combined with various LEE colour filters (https://leefilters.com). Four different light treatments were created: white-cool LED, blue LED, red LED, and green LED. White-cool LED was produced in the same manner as described above. For the blue LED treatment, broad-spectrum LED light (VIPARSPECTRA, Model V165) was set to full intensity (100/100) at the cool LED setting, and the light was passed through a 0.3 neutral density filter (LEE 209), a diffusing sheet, and a blue colour filter (LEE 201). For the green LED treatment, the cool LED intensity was reduced to 70% of its original value, and the light was filtered through a 0.3 neutral density filter, a diffusing sheet, and a green colour filter (LEE 244). The red LED condition was produced by setting the cool LED to 0/100, using a diffusing sheet, and a red colour filter (LEE 106). An additional white-warm LED treatment was produced by using a broad-spectrum white-warm LED light strip without further filtering.

The absolute irradiance in photons/count at 50 lux for each ALAN source was measured using a 1000 μm optical fibre connected to a USB2000 spectrometer (Ocean Optics) on a laptop computer running OceanView software. The fibre was pointed at a 45^0^ angle at a diffusion reflectance standard (WS-1, Ocean Optics) placed at the bottom, in the middle of the experimental section of the tank. This way, the light in the tank was collected from the semisphere above the reflectance standard, representing the overall light intensity experienced in the tank. For each light treatment, the absolute irradiance was measured ten times and averaged (Fig. 3A–E). The light environment during the day, consisting of a mixture of white-warm LED and fluorescent light sources, was measured in the same way (Fig. 3F).

**Figure 3:**
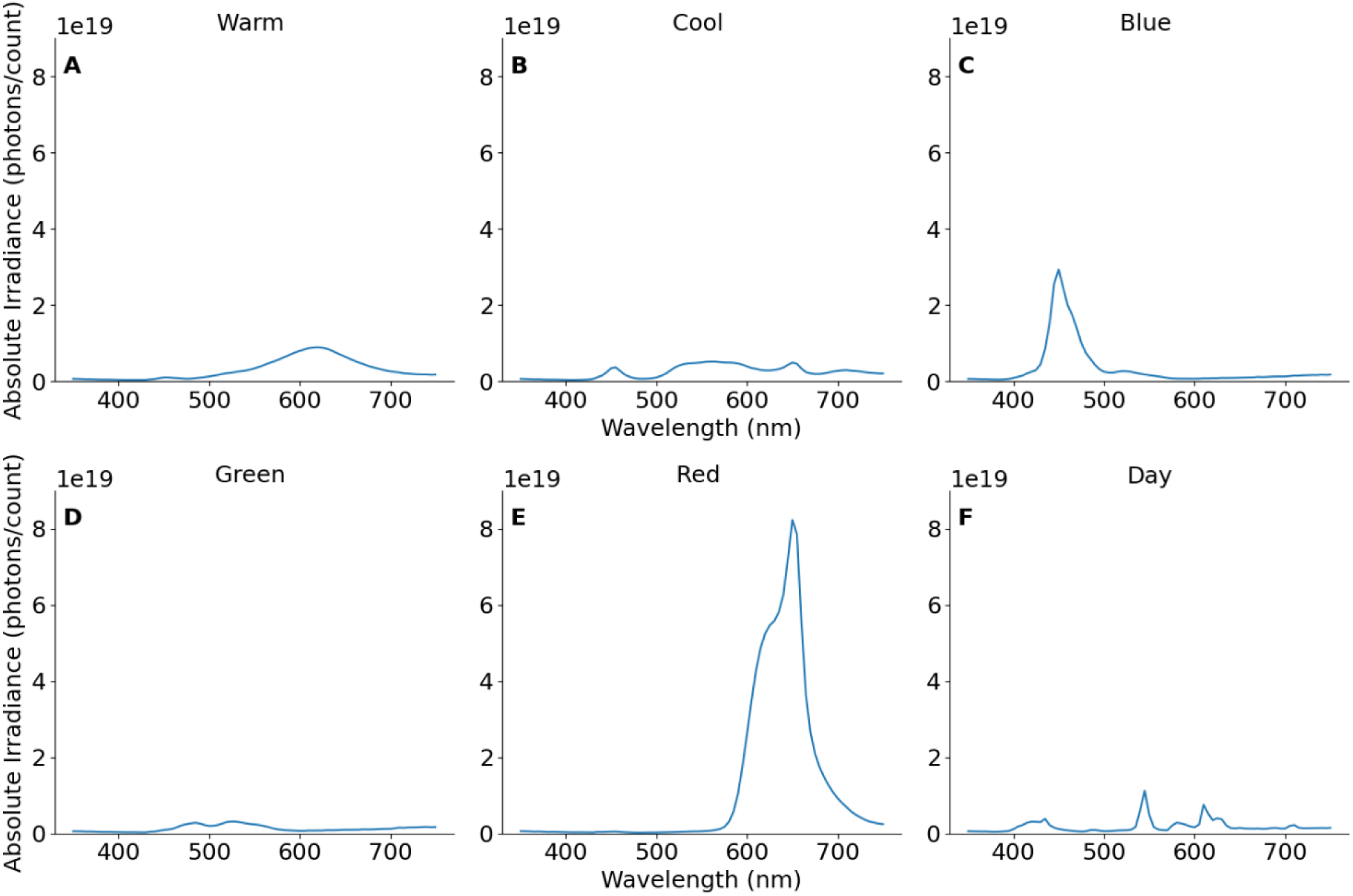
Spectral composition of each ALAN treatment in the discrimination experiment at 50 lux. A – D) ALAN LED-based light sources with: white-warm A), white-cool B), blue C), green D), and red light E). F) Standard laboratory light conditions during the day (a mixture of white-warm LEDs and white-fluorescent lights). Note that, because lux meters are optimised for human vision, at 50 lux, red and blue showed higher light emissions within their respective spectral ranges than the other tested colours. For detailed spectral composition graphs, see Figure S1.

### Statistical analysis of the colour discrimination test

We analysed the binary outcome variable, choice, with 0 indicating failure and 1 indicating success, across multiple trials over several days to assess 1) the effect of different white-cool ALAN intensity and 2) ALAN with different spectral composition on success rates over time. The merged datasets consisted of 2,600 observations from 31 individuals, with repeated measurements per fish.

To account for repeated-measures binomial data, we compared Generalised Linear Mixed Models with a binomial distribution and individual ID as a random factor using the glmer() function from the lme4 package (Boeck et al., 2011) in R v.4.3.3 (Posit Team, 2025). Nested models, including a null model, models with day or treatment alone, an additive model, and a full model with their interactions, were compared using Akaike Information Criterion (AIC). Model selection was based on comparing ΔAIC values and likelihood ratio tests, with the model having ΔAIC < 2 and a non-significant LRT (p > 0.05) chosen as the best-fitting model.

## Results

### The effect of light intensity on memory and learning

Using the white-cool LED treatment at different intensities (0.5, 5 and 50 lux), we first assessed the effect of ALAN light intensity on memory and learning in *A. ocellaris.* For the 0.5 lux treatment, the full interaction model (days × treatment) was selected based on the lowest AIC (ΔAIC = 0), indicating that treatment effects varied over time (Fig. 2A; Tables S1-3). At baseline, success rates were slightly higher in the ALAN treatment (∼88%) than in the control group (∼85%; estimate = 0.30, 95% CI = −0.73 to 1.33). However, while the control group improved over the experiment (∼99% success at the end; estimate = 0.93, 95% CI = 0.27 to 1.59), no improvement was observed in the treatment group (∼87%; interaction estimate = −0.93, 95% CI = −1.74 to −0.12), indicating that learning was impeded in the treatment group. For the 5 lux treatment, the additive model (days + treatment) was selected as the best-supported model, indicating that days and treatment independently influenced success (Fig. 2B; Tables S1-3). The effect of experimental duration was negligible for both groups (estimate = −0.03, 95% CI = −0.38 to 0.32). The control group consistently achieved a very high success rate (∼95%). Conversely, the ALAN treatment group consistently exhibited lower success rates (∼85%; estimate = −1.02, 95% CI = −2.19 to 0.23), indicating that no learning-related improvement occurred in the treatment fish.

In the 50 lux treatment, the interaction model (days × treatment) was selected, suggesting that ALAN modulated the temporal dynamics of success (Fig. 2C; Tables S1-3). At the start of the experiment, treatment fish had a lower success rate than control fish (∼86% vs ∼94%; estimate = −0.98, 95% CI = −2.19 to 0.23). Days had a small positive effect on controls, increasing to ∼98% success at the end (estimate = 0.15, 95% CI = −0.60 to 0.89). Importantly, in the treatment group, 50 lux ALAN reduced success over time (∼69% at the end; interaction estimate = −0.72, 95% CI = −1.56 to 0.12), indicating a strong adverse effect on memory and learning at higher light intensity.

### The effect of spectral composition on memory and learning

The behavioural responses of coral reef fishes to ALAN may be dependent on its spectral composition, as different wavelengths penetrate the water column with varying efficiency and trigger distinct physiological pathways (Holles et al., 2016; Schligler et al., 2021). With ALAN intensity fixed at 50 lux, we assessed the effect of different spectral compositions (Figs. 3 and S1) on memory and learning in *A. ocellaris* (Fig. 4).

**Figure 4:**
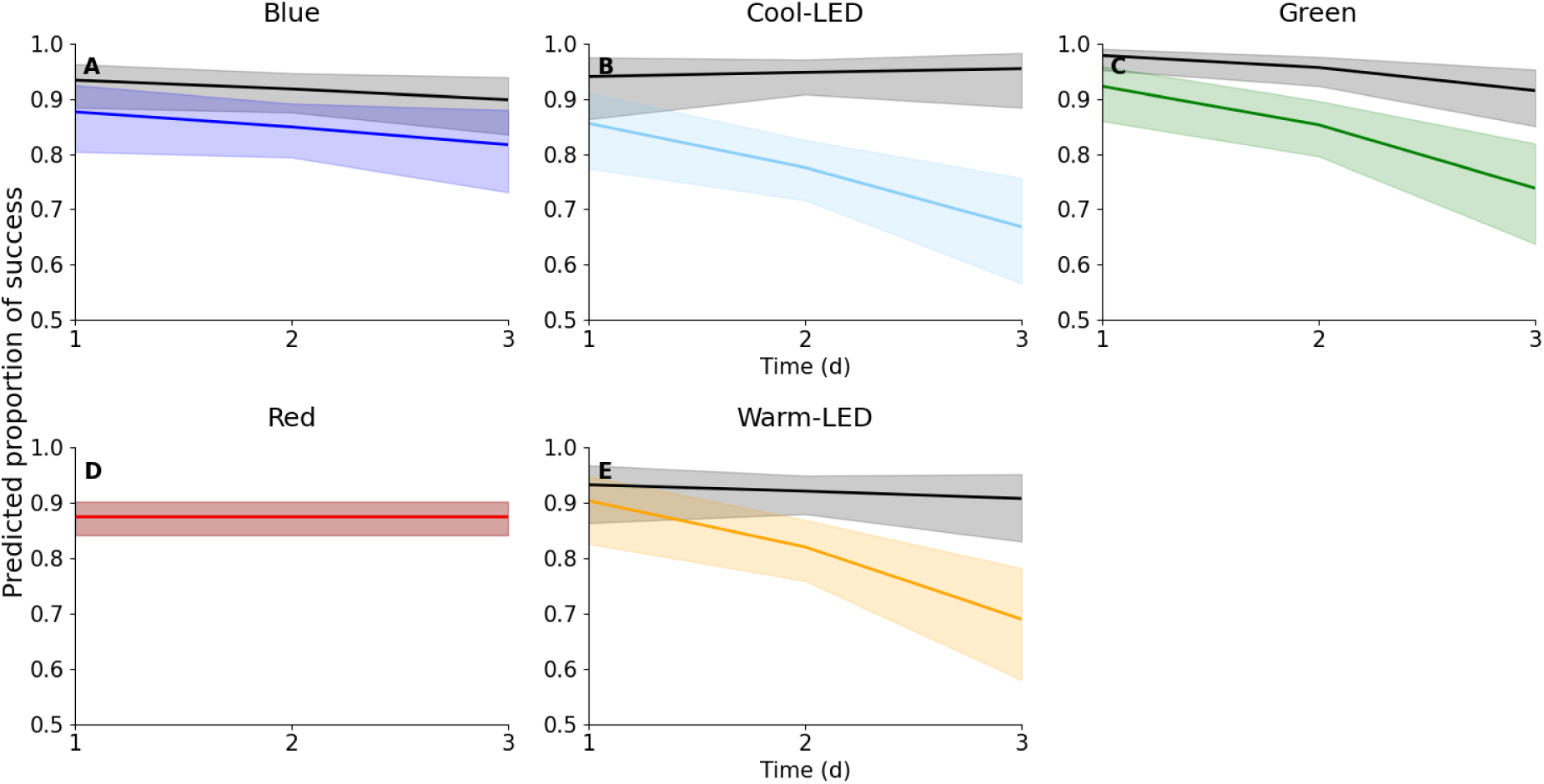
The impact of differently coloured LED lights at night on *Amphiprion ocellaris* memory and learning. Predicted success rates over two consecutive nights of ALAN exposure under 50 lux of blue (A), cool-white LED (B), green (C), red (D), and warm-white LED (E) compared to matched control groups (black) (n = 8 fish per group). Each panel presents the model-predicted mean (solid line) and 95% confidence interval (shaded area). For the red-light treatment, there was no difference between the control and the treatment groups. The y-axis starts at 0.5, representing random chance. See Figs. 3 and S1 for the spectral compositions of the various ALAN treatments.

Using a new cohort of fish, we initially repeated the white-cool LED experiment. As per the findings from the intensity experiment (Fig. 2), the full interaction model (days × treatment) had the highest support (Fig. 4; Tables S4-6). At baseline, success was lower in the white-cool LED treatment (∼86%) than in the control (∼94%; estimate = –0.98, 95% CI = – 2.19 to 0.23). The control group showed modest improvement over the course of the experiment, with a success rate of ∼98% at the end (estimate = 0.15, 95% CI = –0.60 to 0.89). Conversely, the white-cool LED treatment greatly reduced success over time (∼68%; interaction estimate = –0.72, 95% CI = –1.56 to 0.12), repeating the results from the intensity experiment (Figs. 3C, 4B).

Based on the strong adverse effects on learning and memory observed with white-cool LEDs, we then tested the impact of white-warm LED lights. These lights showed reduced emission in the blue-green spectrum and increased emission in the yellow-red range compared to white-cool LEDs (Fig. 3A). The full interaction model (days × treatment) was again selected (Fig. 4; Tables S4-6). At the start of the experiment, treatment fish (∼91%) had a slightly lower success rate than controls (∼94%; estimate = –0.38, 95% CI = –1.47 to 0.71). Days had little effect on control, with success rates staying consistently high (∼95%; estimate = –0.17, 95% CI = –0.75 to 0.40). However, the interaction between days and treatment showed a substantial reduction in success over time in the white-warm LED treatment, with a success rate of ∼70% at the end of the experiment (interaction estimate = –0.56, 95% CI = –1.30 to 0.18). Hence, both white-LED treatments greatly reduced memory and learning in *A. ocellaris*, with white-cool LEDs showing a slightly more pronounced effect (Fig. 4B, E).

Next, we tested the separate spectral ranges that make up human-visible white light. Starting from the shorter wavelength end, we first tested the effects of blue and green LEDs, as they penetrate the water the furthest (Jerlov, 1976). For blue LEDs, the highest support was for the additive model (days + treatment), while for green LEDs, the highest support was for the full interaction model (days × treatment) (Fig. 4, Tables S4-6). In the blue LED experiment, success rates at baseline were similar between the treatment and control groups (90-95%; estimate = 2.37, 95% CI = 1.80 to 2.94). The level of success for both groups dropped slightly over time (estimate = –0.23, 95% CI = –0.59 to 0.12). However, while the control group maintained a success rate well above 90%, the treatment group dropped to just below 85% (Fig. 4A), which we considered the minimum level of success beyond random chance for our experiments (see Methods above).

For the green LED, at the start of the experiment, success, albeit high overall, was lower in the treatment group (∼92%) compared to the control (∼99%; estimate = –0.99, 95% CI = – 2.46 to 0.47). The success of the control dropped slightly, to ∼97% at the end of the experiment (estimate = –0.55, 95% CI = –1.31 to 0.20). Conversely, the success rate of the green ALAN treatment dropped to ∼78% at the end of the experiment (estimate = –0.26, 95% CI = –1.17 to 0.65). Hence, both green and blue LED lights reduced memory and learning in *A. ocellaris* to levels well below the random-choice level, or just at that level, respectively (Fig. 4A, C).

Finally, we tested the effects of red LED ALAN on cognitive performance in the fish, as this spectral range showed the largest difference between white-cool and white-warm LED lights (Fig. 3). Interestingly, despite being much brighter than the other treatment groups, the best model fit was found for the null model that did not include days and treatment. This means that the success rate was similar between groups at the start of the experiment (estimate = 2.13, 95% CI = 1.57 to 2.69), and it remained at ∼89% throughout the entire experiment (estimate = –0.03, 95% CI = –0.36 to 0.30) (Fig. 4D; Tables S4-6). This suggests that, while broad-spectrum or shorter-spectrum LEDs have an adverse effect on *A. ocellaris* cognition, the fish appear indifferent to red light at night.

## Discussion

The effects of ALAN on reef fish cognition and behaviour are poorly understood (Ferretti et al., 2024; Weschke, 2024). In this study, we aimed to investigate whether different intensities and colours of ALAN impact memory in the clownfish, *A. ocellaris*. Clownfishes are representative of the many smaller, site-attached fishes on shallow tropical coral reefs, which experience a rapid increase in ALAN in their natural environment (Laudet & Ravasi, 2023). Testing contemporary white-cool LED lights at different intensities revealed that even at very low light levels of 0.5 lux, *A. ocellaris* learning was impeded (Fig. 2A). At the highest level of 50 lux tested here, not only learning, but memory itself was clearly affected, with fish mostly resorting to random choice when looking for their food reward (Fig. 2C). Supporting the importance of shorter-wavelengths of light for the visual ecology of diurnal reef fishes (Cortesi et al., 2020), broad-spectrum white LEDs (warm and cool) and green and blue LEDs all negatively impacted learning and memory in *A. ocellaris*. Red LEDs, on the other hand, did not affect fish cognition, even when presented at a higher intensity than the other colour treatments (Fig. 4).

Memory and learning are essential for every aspect of a fish’s ecology, including migration, foraging, landmark orientation, social behaviours and adaptation to environmental change (Kieffer & Colgan, 1992; Munson & DePasquale, 2025; Overmier & Hollis, 1990). While the largest effects were observed for street-light intensities (50 lux), even low levels of ALAN at full moon intensities (0.5 lux) had an impact on learning in *A. ocellaris* (Fig. 2). Such low levels of ALAN have the potential to be far spread, especially in areas with indirect illumination via skyglow, and should be focused on more thoroughly in the future (Smyth et al., 2021; Weschke, 2024). Moreover, both chronic ALAN exposure, as found in many urbanised coastal areas (Davies et al., 2016), and the natural light fluctuations at night over multiple lunar cycles (Tidau et al., 2021), should be studied to determine if low levels of ALAN cause additional stress, or whether these effects fall within the range that coral reef fishes naturally experience.

Retinal cone spectral sensitivities in *A. ocellaris* peak at 386 (ultraviolet-violet), 497 (blue-green), 515 (green), and 535 (green-yellow) nm λ_max_ (Mitchell et al., 2024). The species also uses a single rod photoreceptor with a λ_max_ at 491–499 nm for scotopic vision (Mitchell et al., 2021). These sensitivity peaks align most closely with the spectral compositions of the white-cool-, white-warm-, and green LED treatments used in our study (Figs. 3 and S1). The broader spectral output of the white LEDs likely enabled greater overlap with multiple photoreceptor sensitivities, enhancing their detectability by *A. ocellaris* (Fig. 4). The two peaks in the green LED output (Fig. S1) overlap almost perfectly with the two mid-wavelength sensitive cone photoreceptors (λ_max_ at 497 and 515 nm) as well as the spectral sensitivity of the rod photoreceptor (Mitchell et al., 2024). Accordingly, the green treatment alone was enough to replicate most of the effect found for the broadband white-light treatments (Fig. 4). As green had the lowest intensity of all the tested light treatments (Fig. S1), increasing its intensity to match the maximum output in the other treatments would likely have caused an even stronger effect.

In contrast, despite being 2–10 times brighter at peak emission (Fig. S1), neither blue nor red LEDs had a strong effect on *A. ocellaris* performance. This might be explained by the fact that the LED outputs (Fig. S1) only partially overlap with the photoreceptor spectral sensitivities of *A. ocellaris* (Mitchell et al., 2024). Hence, they might not be able to perceive these lights well. Indeed, when testing the colour discrimination ability of this species, Mitchell and colleagues found that they do not perform well in the blue part of the spectrum (Mitchell et al., 2024). However, as far as red colour is concerned, *A. ocellaris* has high contrast sensitivity in this part of the spectrum that matches or even outperforms that of the green spectrum (Mitchell et al., 2024). This is likely due to their algal food reflecting strongly in the far-red; clownfishes also use orange and red colours for signalling, and because their symbiotic anemones often have a red colouration (Stieb et al., 2023). It is unclear whether a longer, chronic exposure to red light would eventually have impaired *A. ocellaris* cognition. Either way, the indifference to the red LED ALAN we discovered here is somewhat perplexing, and the underlying neurophysiological principles warrant further investigation. As such, it is plausible that the cone photoreceptors in the eye are not the primary light sensors involved in perceiving ALAN, as the retinal epithelium movement (black pigment layer) likely protects them from light exposure (Ali, 1975; Burnside, 2001). Instead, both the green-sensitive rods in the eye (Mitchell et al., 2021) and the putatively green-sensitive pineal gland photoreceptors might be the primary sensors contributing to the behavioural disturbances observed. Both are central to the circadian rhythm, which regulates sleep in fishes, with sleep disturbance and ALAN being irreversibly tied to one another (Sigurgeirsson et al., 2013). Hence, future work assessing sleep in the clownfish, combined with molecular analyses of retinal and pineal gland gene expression, may elucidate the mechanisms underlying the learning and memory impairments observed in this study.

In general, many shallow-living reef fish species, and also many pelagic species, have visual systems that are similar to the visual system of *A. ocellaris* (reviewed in Marshall et al., 2019; Cortesi et al., 2020). This suggests that broadband white LEDs, especially those using the newer white-cool LEDs, are likely to have a widespread adverse effect on the reef fish community. Additionally, because many reef fish species utilise a distinct blue cone photoreceptor (Cortesi et al., 2020), which is missing in *A. ocellaris*, blue-only light sources are likely to have a more substantial effect than observed here. At first glance, then, focusing management efforts on using longer-wavelength emitting red light sources might be the best solution. However, different marine species have shown contrasting reactions to differently coloured light sources. For example, blue light accelerates development and metamorphosis in marine shellfish larvae (Zhang et al., 2023) and reduces the bycatch of fish and sea turtles (Stanton & Cowart, 2024). Contrarily, blue light impairs the locomotion of gastropod grazers (Maggi et al., 2020). Red and green lights are considered less disruptive to turtle nesting and hatchling orientation (Cruz et al., 2018), North Atlantic mesopelagic fish (Peña et al., 2020), and seabird navigation (Syposz et al., 2021). Yet, these same wavelengths disrupt Arctic pelagic organisms (Geoffroy et al., 2021) and increase the biomass of primary producers, leading to the selective dominance of specific diatom species known to cause harmful algal blooms (Diamantopoulou et al., 2021). As shown here, especially using green light at night, also disrupts learning and memory in clownfish (Fig. 4). Therefore, future efforts should focus on investigating whether the indifference to longer-wavelength ALAN observed in the clownfish is an isolated case or whether it is widespread in the reef (fish) community.

## Conclusion

Our study contributes to a growing body of literature on the effects of ALAN on the cognitive abilities of reef fish. The finding that *A. ocellaris* is indifferent to red light at night adds an essential aspect to the broader discussion of marine ALAN management. It suggests that replacing broadband LEDs with longer-wavelength red LEDs may minimise effects on shallow living coral reef fishes. However, because species differ in their responses to the same spectrum/colour of light, it is unlikely that any single “neutral” light source can minimise ecological impacts across all taxa. Moreover, our finding that learning and memory in clownfish were dependent on the intensity of ALAN used reinforces the general recommendation for the design of light sources at night in urban and coastal planning: reducing overall ALAN exposure, or, as a bare minimum, its intensity, remains the most effective and ecologically responsible strategy to protect the marine environment.

## Data Availability

All the data is available from the Supplementary Materials and Methods or upon request from the corresponding author.

## Acknowledgements

We thank Angelo Guadagno and Dr Zegni Triki for their assistance with the behavioural assay and their patience in teaching us how to implement it. We also thank the members of the Marine Sensory Ecology group for their help with aquarium husbandry and for their inspiration. This work was funded by an Australian Research Council Future Fellowship (FT240100725) to FC.

## Author Contributions

F.C. designed the experiments. J.vST. and C.M. performed the experiments, and J.vST. analysed the data. J.vSt. and F.C. wrote the initial version, and all authors have contributed to the final version of the manuscript.

## Competing interests

The authors declare no competing interests.

## Supplementary Materials and Methods

**Figure S1:**
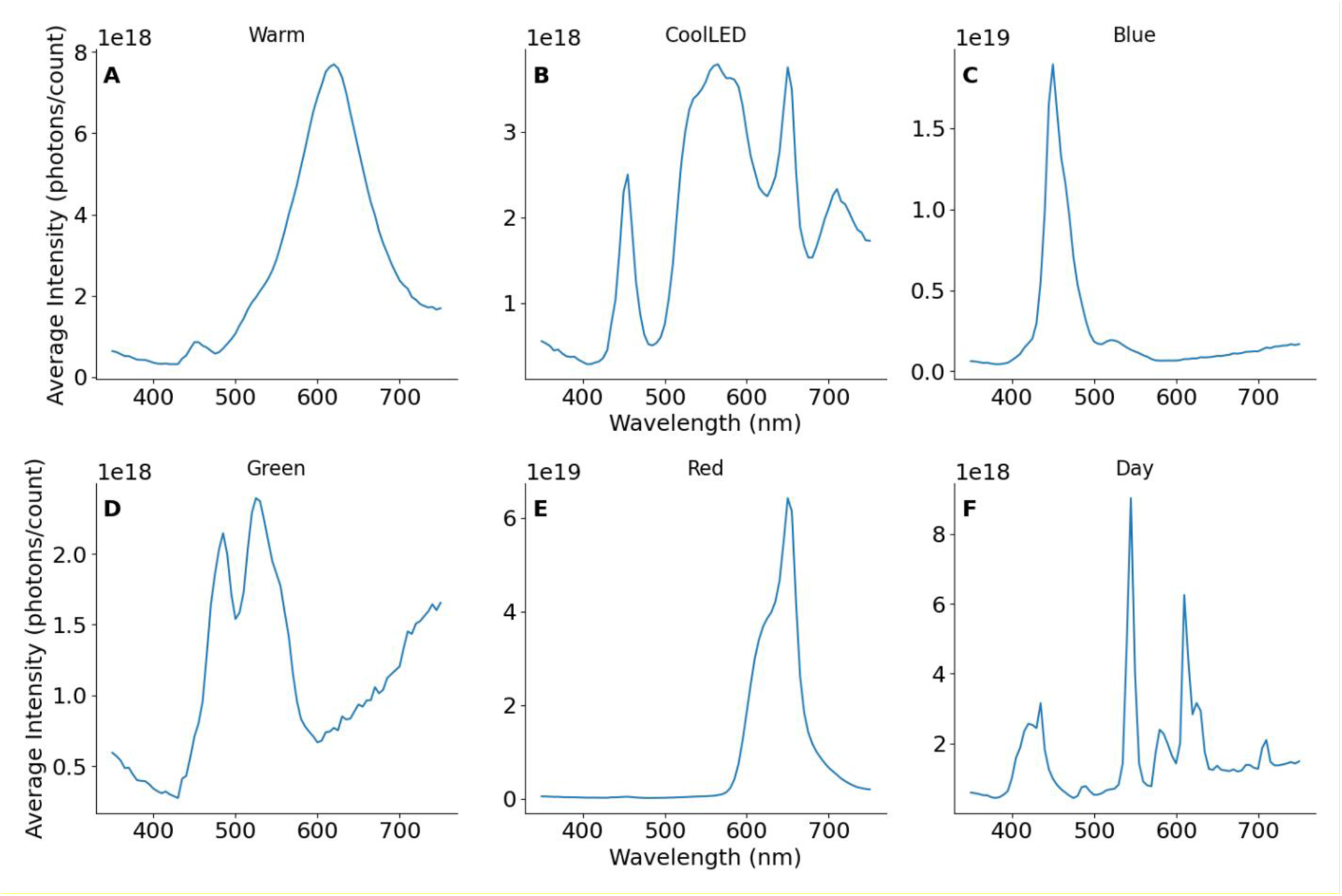
Absolute irradiance (photons/count) at 50 lux, and spectral composition of each light treatment in the differently coloured LED experiment, measured at the centre of the tank (averaged across 10 measurements). White-warm LED (A), white-cool LED (B), blue LED (C), green LED (D), red LED (E), standard laboratory light conditions during the day (F).

**Table S1:** Akaike Information Criterion (AIC) comparison of Generalised Linear Mixed Models analysing *Amphiprion ocellaris’* success in the light intensity experiment. AIC = Akaike Information Criterion; Model = model ID; ΔAIC = difference in AIC relative to the best-supported model. Models: m0 = Success ∼ 1 + (1 | Fish), m1 = Success ∼ Days + (1 | Fish), m2 = Success ∼ Days + Treatment + (1 | Fish), m3 = Success ∼ Days * Treatment + (1 | Fish).

| Degrees of freedom | AIC | Model | Treatment group | ΔAIC |
| --- | --- | --- | --- | --- |
| 2 | 318.87 | m0 | 0.5 lux | 3.64 |
| 3 | 317.36 | m1 | 0.5 lux | 2.13 |
| 4 | 318.49 | m2 | 0.5 lux | 3.26 |
| 5 | 315.22 | m3 | 0.5 lux | 0 |
| 2 | 333.57 | m0 | 5 lux | 1.11 |
| 3 | 335.54 | m1 | 5 lux | 3.08 |
| 4 | 332.46 | m2 | 5 lux | 0 |
| 5 | 333.64 | m3 | 5 lux | 1.18 |
| 2 | 358.05 | m0 | 50 lux | 14.60 |
| 3 | 354.21 | m1 | 50 lux | 10.77 |
| 4 | 344.18 | m2 | 50 lux | 0.74 |
| 5 | 343.44 | m3 | 50 lux | 0 |

**Table S2:** Likelihood Ratio Test (LRT) comparison of Generalised Linear Mixed Models analysing A. occelaris’ success in the 0.5, 5, and 50 lux treatments. Term = model; AIC = Akaike Information Criterion; BIC = Bayesian Information Criterion; statistic = LRT test statistic. Models: m0 = Success ∼ 1 + (1 | Fish), m1 = Success ∼ Days + (1 | Fish), m2 = Success ∼ Days + Treatment + (1 | Fish), m3 = Success ∼ Days * Treatment + (1 | Fish).

| Term | Number of parameters | AIC | BIC | Log-likelihood | -2 × log-likelihood | Statistic | Degrees of freedom | p-value | Treatment group |
| --- | --- | --- | --- | --- | --- | --- | --- | --- | --- |
| m0 | 2 | 318.87 | 327.22 | -157.43 | 314.87 |  |  |  | Pair_0_5 |
| m1 | 3 | 317.36 | 329.88 | -155.68 | 311.36 | 3.50 | 1 | 0.06 | Pair_0_5 |
| m2 | 4 | 318.49 | 335.19 | -155.24 | 310.49 | 0.86 | 1 | 0.35 | Pair_0_5 |
| m3 | 5 | 315.22 | 336.09 | -152.61 | 305.22 | 5.26 | 1 | 0.02 | Pair_0_5 |
| m0 | 2 | 333.57 | 341.92 | -164.78 | 329.57 |  |  |  | Pair_5 |
| m1 | 3 | 335.54 | 348.06 | -164.77 | 329.54 | 0.03 | 1 | 0.85 | Pair_5 |
| m2 | 4 | 332.46 | 349.15 | -162.23 | 324.46 | 5.08 | 1 | 0.02 | Pair_5 |
| m3 | 5 | 333.64 | 354.51 | -161.82 | 323.64 | 0.81 | 1 | 0.36 | Pair_5 |
| m0 | 2 | 358.05 | 366.26 | -177.02 | 354.05 |  |  |  | Pair_50 |
| m1 | 3 | 354.21 | 366.54 | -174.10 | 348.21 | 5.83 | 1 | 0.01 | Pair_50 |
| m2 | 4 | 344.18 | 360.62 | -168.09 | 336.18 | 12.02 | 1 | 0.0005 | Pair_50 |
| m3 | 5 | 343.44 | 363.98 | -166.72 | 333.44 | 2.74 | 1 | 0.09 | Pair_50 |

**Table S3:** Parameter estimates and 95% confidence intervals for the best-supported Generalised Linear Mixed Models predicting A. occelaris’ success at 0.5, 5, and 50 lux. Model = selected model; Formula = model formula; Selection reason = rationale for model selection; term= model term; estimate = model coefficient; lower 95%CI/upper 95% CI = lower and upper 95% confidence intervals; effect = fixed or random effect; group = group assignment; SE = standard error; statistic = test statistic.

| Treatment group | Model | Formula | Selection Reason | Term | Estimate | Lower 95% CI | Upper 95% CI | Effect | Group | SE | Statistic | P-value |
| --- | --- | --- | --- | --- | --- | --- | --- | --- | --- | --- | --- | --- |
| 0.5 lux | m3 | Success ~ Days * Treatment + (1 \| Fish) | Lowest AIC | (Intercept) | 1.79 | 1.07 | 2.51 | fixed |  | 0.36 | 4.87 | 1.1E-06 |
| 0.5 lux | m3 | Success ~ Days * Treatment + (1 \| Fish) | Lowest AIC | Days | 0.92 | 0.26 | 1.58 | fixed |  | 0.33 | 2.76 | 0.005 |
| 0.5 lux | m3 | Success ~ Days * Treatment + (1 \| Fish) | Lowest AIC | Treatment | 0.29 | -0.73 | 1.33 | fixed |  | 0.52 | 0.56 | 0.57 |
| 0.5 lux | m3 | Success ~ Days * Treatment + (1 \| Fish) | Lowest AIC | Days: Treatment | -0.92 | -1.73 | -0.11 | fixed |  | 0.41 | -2.24 | 0.02 |
| 0.5 lux | m3 | Success ~ Days * Treatment + (1 \| Fish) | Lowest AIC | SD (Intercept) | 0.57 |  |  | random | Fish |  |  |  |
| 5 lux | m1 | Success ~ Days + (1 \| Fish) | $\Delta AIC < 2$ and LRT p > 0.05 | (Intercept) | 2.31 | 1.68 | 2.94 | fixed | | 0.32 | 7.15 | 8.39E-13 |
| 5 lux | m1 | Success ~ Days + (1 \| Fish) | $\Delta AIC < 2$ and LRT p > 0.05 | Days | -0.03 | -0.38 | 0.31 | fixed | | 0.17 | -0.18 | 0.85 |
| 5 lux | m1 | Success ~ Days + (1 \| Fish) | $\Delta AIC < 2$ and LRT p > 0.05 | SD (Intercept) | 0.77 | | | random | Fish | | | |
| 50 lux | m3 | Success ~ Days * Treatment + (1 \| Fish) | $\Delta AIC < 2$ and LRT p > 0.05 | (Intercept) | 2.87 | 1.86 | 3.88 | fixed | | 0.51 | 5.56 | 2.6E-08 |
| 50 lux | m3 | Success ~ Days * Treatment + (1 \| Fish) | $\Delta AIC < 2$ and LRT p > 0.05 | Days | 0.14 | -0.59 | 0.88 | fixed | | 0.37 | 0.38 | 0.69 |
| 50 lux | m3 | Success ~<br>Days *<br>Treatment + (1<br>\| Fish) | $\Delta AIC < 2$<br>and LRT p<br>> 0.05 | Treatment | -0.97 | -2.18 | 0.23 | fixed | | 0.61 | -1.58 | 0.11 |
| 50 lux | m3 | Success ~<br>Days *<br>Treatment + (1<br>\| Fish) | $\Delta AIC < 2$<br>and LRT p<br>> 0.05 | Days:<br>Treatment | -0.71 | -1.55 | 0.12 | fixed | | 0.42 | -1.67 | 0.09 |
| 50 lux | m3 | Success ~<br>Days *<br>Treatment + (1<br>\| Fish) | $\Delta AIC < 2$<br>and LRT p<br>> 0.05 | SD<br>(intercept) | 0.54 | | | random | Fish | | | |

**Table S4:** Akaike Information Criterion (AIC) comparison of Generalised Linear Mixed Models analysing A. occelaris’ success under various spectral compositions of ALAN treatment. AIC = Akaike Information Criterion; Model = model ID; ΔAIC = difference in AIC relative to the best-supported model. Models: m0 = Success ∼ 1 + (1 | Fish), m1 = Success ∼ Days + (1 | Fish), m2 = Success ∼ Days + Treatment + (1 | Fish), m3 = Success ∼ Days * Treatment + (1 | Fish).

| Degrees of freedom | AIC | Model | Treatment group | ΔAIC |
| --- | --- | --- | --- | --- |
| 2 | 358.05 | m0 | Cool LED | 14.60 |
| 3 | 354.21 | m1 | Cool LED | 10.77 |
| 4 | 344.18 | m2 | Cool LED | 0.74 |
| 5 | 343.44 | m3 | Cool LED | 0 |
| 2 | 324.27 | m0 | Blue LED | 2.59 |
| 3 | 324.63 | m1 | Blue LED | 2.95 |
| 4 | 322.97 | m2 | Blue LED | 1.29 |
| 5 | 321.67 | m3 | Blue LED | 0 |
| 2 | 358.20 | m0 | Red LED | 0 |
| 3 | 360.17 | m1 | Red LED | 1.97 |
| 4 | 361.24 | m2 | Red LED | 3.04 |
| 5 | 362.94 | m3 | Red LED | 4.73 |
| 2 | 292.11 | m0 | Green LED | 17.06 |
| 3 | 281.28 | m1 | Green LED | 6.23 |
| 4 | 275.04 | m2 | Green LED | 0 |
| 5 | 276.74 | m3 | Green LED | 1.69 |
| 2 | 351.47 | m0 | Warm LED | 12.95 |
| 3 | 344.65 | m1 | Warm LED | 6.12 |
| 4 | 338.72 | m2 | Warm LED | 0.19 |
| 5 | 338.52 | m3 | Warm LED | 0 |

**Table S5:** Likelihood Ratio Test (LRT) comparison of Generalised Linear Mixed Models analysing A. occelaris’ success under various spectral compositions of ALAN treatment. Term = model; AIC = Akaike Information Criterion; BIC = Bayesian Information Criterion; statistic = LRT test statistic. Models: m0 = Success ∼ 1 + (1 | Fish), m1 = Success ∼ Days + (1 | Fish), m2 = Success ∼ Days + Treatment + (1 | Fish), m3 = Success ∼ Days * Treatment + (1 | Fish).

| Term | Number of parameters | AIC | BIC | log-likelihood | -2 × log-likelihood | statistic | Degrees of freedom | p-value | Treatment group |
| --- | --- | --- | --- | --- | --- | --- | --- | --- | --- |
| m0 | 2 | 358.05 | 366.26 | -177.02 | 354.05 |  |  |  | Cool LED |
| m1 | 3 | 354.21 | 366.54 | -174.10 | 348.21 | 5.83 | 1 | 0.015 | Cool LED |
| m2 | 4 | 344.18 | 360.62 | -168.09 | 336.18 | 12.02 | 1 | 0.0005 | Cool LED |
| m3 | 5 | 343.44 | 363.98 | -166.72 | 333.44 | 2.74 | 1 | 0.09 | Cool LED |
| m0 | 2 | 324.27 | 332.49 | -160.13 | 320.27 |  |  |  | Blue LED |
| m1 | 3 | 324.63 | 336.96 | -159.31 | 318.63 | 1.63 | 1 | 0.20 | Blue LED |
| m2 | 4 | 322.97 | 339.40 | -157.48 | 314.97 | 3.66 | 1 | 0.05 | Blue LED |
| m3 | 5 | 321.67 | 342.22 | -155.83 | 311.67 | 3.29 | 1 | 0.06 | Blue LED |
| m0 | 2 | 358.20 | 366.55 | -177.10 | 354.20 |  |  |  | Red LED |
| m1 | 3 | 360.17 | 372.69 | -177.08 | 354.17 | 0.02 | 1 | 0.86 | Red LED |
| m2 | 4 | 361.24 | 377.94 | -176.62 | 353.24 | 0.92 | 1 | 0.33 | Red LED |
| m3 | 5 | 362.94 | 383.81 | -176.47 | 352.94 | 0.30 | 1 | 0.58 | Red LED |
| m0 | 2 | 292.11 | 300.33 | -144.05 | 288.11 |  |  |  | Green LED |
| m1 | 3 | 281.28 | 293.60 | -137.64 | 275.28 | 12.83 | 1 | 0.0003 | Green LED |
| m2 | 4 | 275.04 | 291.48 | -133.52 | 267.04 | 8.23 | 1 | 0.004 | Green LED |
| m3 | 5 | 276.74 | 297.28 | -133.37 | 266.74 | 0.30 | 1 | 0.58 | Green LED |
| m0 | 2 | 351.47 | 359.69 | -173.74 | 347.47 |  |  |  | Warm LED |
| m1 | 3 | 344.65 | 356.98 | -169.32 | 338.65 | 8.82 | 1 | 0.002 | Warm LED |
| m2 | 4 | 338.72 | 355.16 | -165.36 | 330.72 | 7.93 | 1 | 0.004 | Warm LED |
| m3 | 5 | 338.52 | 359.07 | -164.26 | 328.52 | 2.19 | 1 | 0.13 | Warm LED |

**Table S6:**
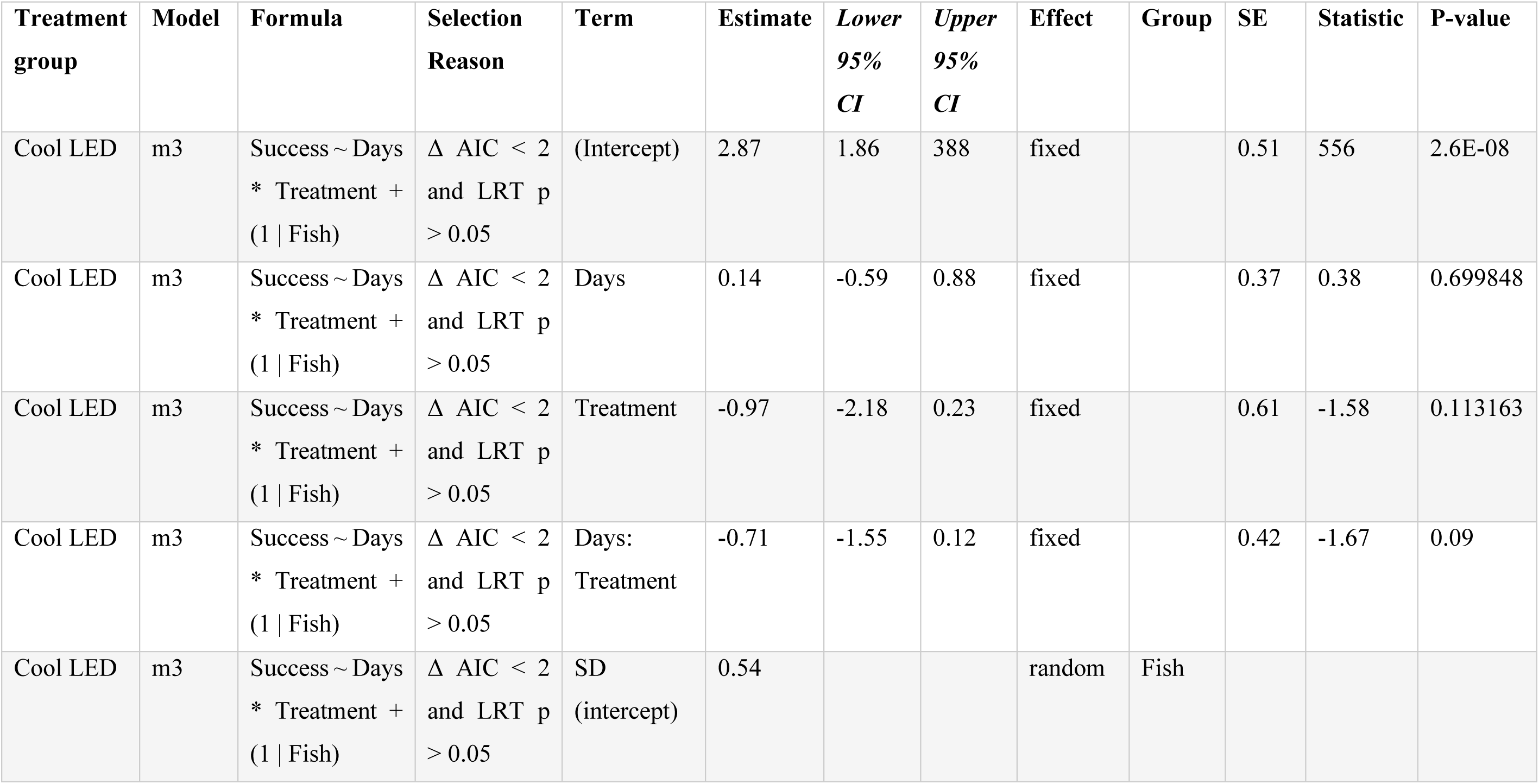
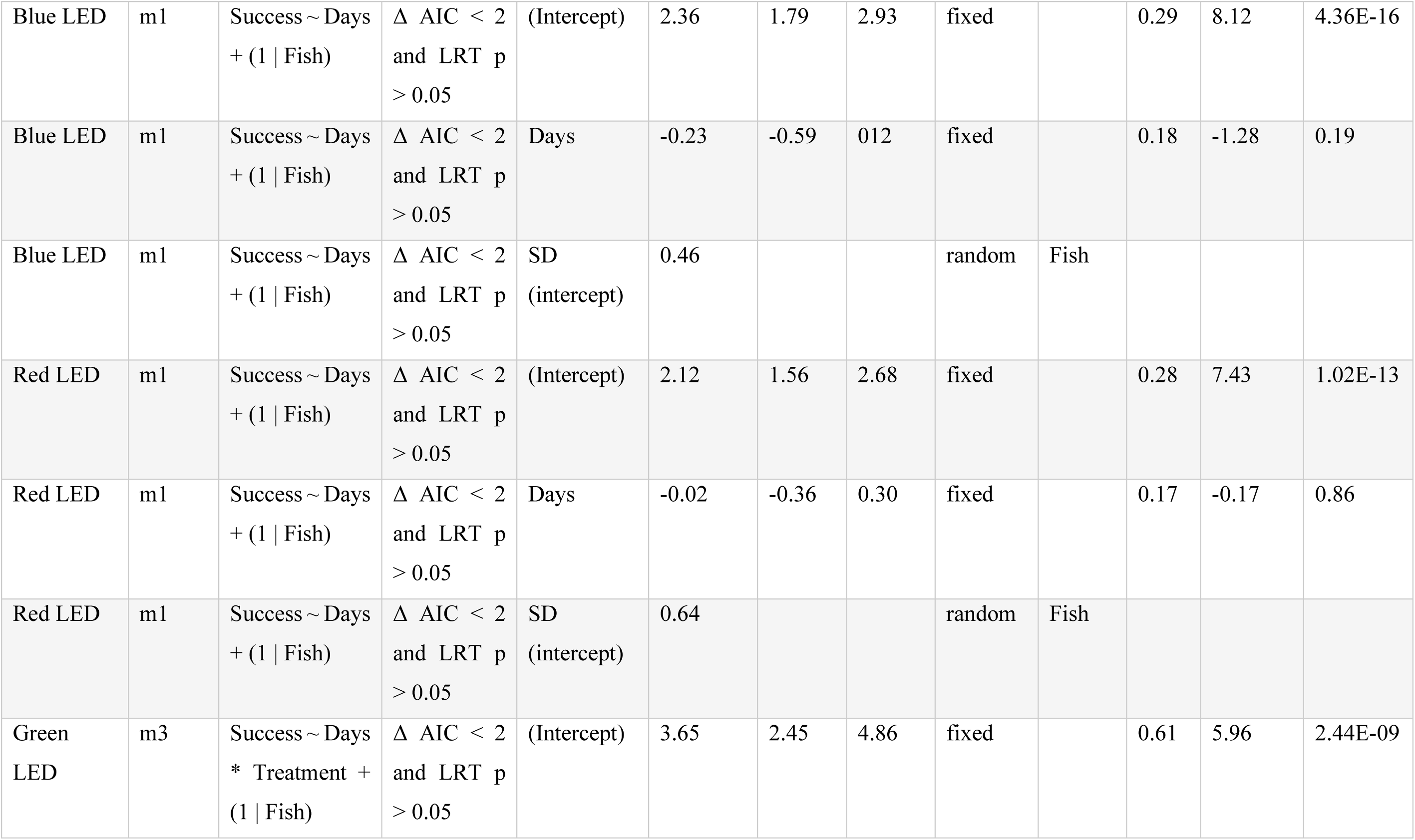
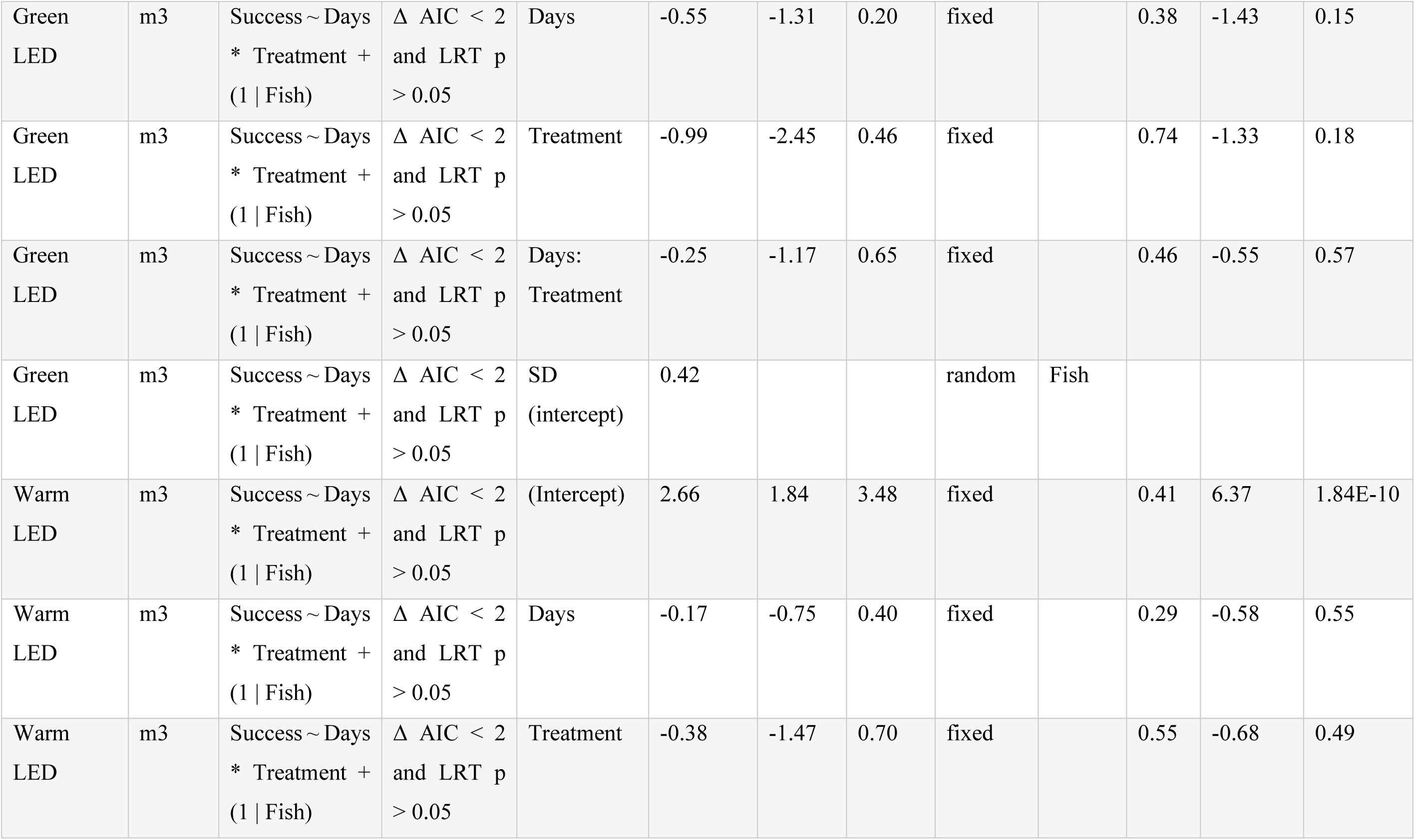
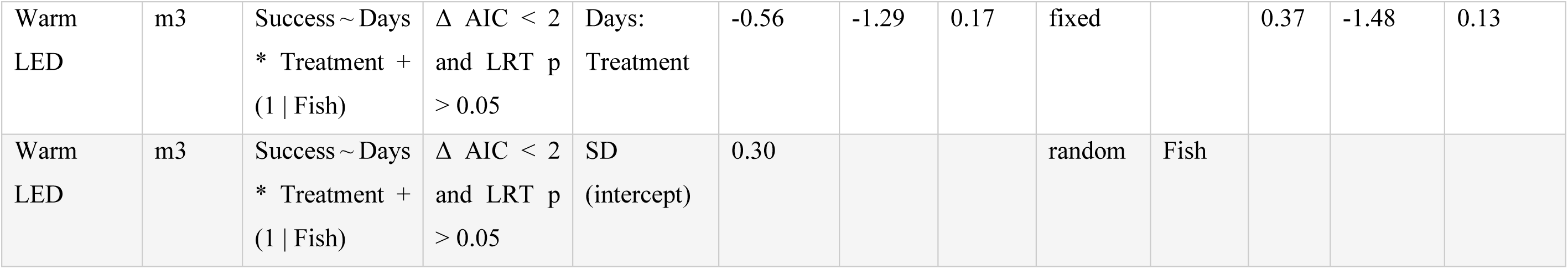
Parameter estimates and 95% confidence intervals for the best-supported Generalised Linear Mixed Models predicting A. occelaris’ success at 0.5, 5, and 50 lux. Treatment group = ALAN treatment; Model = selected model; Formula = model formula; Selection Reason = rationale for model selection; term= model term; estimate = model coefficient; lower 95%CI/upper 95% CI = lower and upper 95% confidence intervals; effect = fixed or random effect; group = group assignment; SE = standard error; statistic = test statistic.

| Treatment group | Model | Formula | Selection Reason | Term | Estimate | Lower 95% CI | Upper 95% CI | Effect | Group | SE | Statistic | P-value |
| --- | --- | --- | --- | --- | --- | --- | --- | --- | --- | --- | --- | --- |
| Cool LED | m3 | Success ~ Days<br>* Treatment +<br>(1 \| Fish) | $\Delta$ AIC < 2<br>and LRT p<br>> 0.05 | (Intercept) | 2.87 | 1.86 | 388 | fixed | | 0.51 | 556 | 2.6E-08 |
| Cool LED | m3 | Success ~ Days<br>* Treatment +<br>(1 \| Fish) | $\Delta$ AIC < 2<br>and LRT p<br>> 0.05 | Days | 0.14 | -0.59 | 0.88 | fixed | | 0.37 | 0.38 | 0.699848 |
| Cool LED | m3 | Success ~ Days<br>* Treatment +<br>(1 \| Fish) | $\Delta$ AIC < 2<br>and LRT p<br>> 0.05 | Treatment | -0.97 | -2.18 | 0.23 | fixed | | 0.61 | -1.58 | 0.113163 |
| Cool LED | m3 | Success ~ Days<br>* Treatment +<br>(1 \| Fish) | $\Delta$ AIC < 2<br>and LRT p<br>> 0.05 | Days:<br>Treatment | -0.71 | -1.55 | 0.12 | fixed | | 0.42 | -1.67 | 0.09 |
| Cool LED | m3 | Success ~ Days<br>* Treatment +<br>(1 \| Fish) | $\Delta$ AIC < 2<br>and LRT p<br>> 0.05 | SD<br>(intercept) | 0.54 | | | random | Fish | | | |
| Blue LED | m1 | Success ~ Days<br>+ (1 \| Fish) | $\Delta$ AIC < 2<br>and LRT p<br>> 0.05 | (Intercept) | 2.36 | 1.79 | 2.93 | fixed | | 0.29 | 8.12 | 4.36E-16 |
| Blue LED | m1 | Success ~ Days<br>+ (1 \| Fish) | $\Delta$ AIC < 2<br>and LRT p<br>> 0.05 | Days | -0.23 | -0.59 | 0.12 | fixed | | 0.18 | -1.28 | 0.19 |
| Blue LED | m1 | Success ~ Days<br>+ (1 \| Fish) | $\Delta$ AIC < 2<br>and LRT p<br>> 0.05 | SD<br>(intercept) | 0.46 | | | random | Fish | | | |
| Red LED | m1 | Success ~ Days<br>+ (1 \| Fish) | $\Delta$ AIC < 2<br>and LRT p<br>> 0.05 | (Intercept) | 2.12 | 1.56 | 2.68 | fixed | | 0.28 | 7.43 | 1.02E-13 |
| Red LED | m1 | Success ~ Days<br>+ (1 \| Fish) | $\Delta$ AIC < 2<br>and LRT p<br>> 0.05 | Days | -0.02 | -0.36 | 0.30 | fixed | | 0.17 | -0.17 | 0.86 |
| Red LED | m1 | Success ~ Days<br>+ (1 \| Fish) | $\Delta$ AIC < 2<br>and LRT p<br>> 0.05 | SD<br>(intercept) | 0.64 | | | random | Fish | | | |
| Green<br>LED | m3 | Success ~ Days<br>* Treatment +<br>(1 \| Fish) | $\Delta$ AIC < 2<br>and LRT p<br>> 0.05 | (Intercept) | 3.65 | 2.45 | 4.86 | fixed | | 0.61 | 5.96 | 2.44E-09 |
| Green LED | m3 | Success ~ Days<br>* Treatment +<br>(1 \| Fish) | $\Delta AIC < 2$<br>and LRT p<br>> 0.05 | Days | -0.55 | -1.31 | 0.20 | fixed | | 0.38 | -1.43 | 0.15 |
| Green LED | m3 | Success ~ Days<br>* Treatment +<br>(1 \| Fish) | $\Delta AIC < 2$<br>and LRT p<br>> 0.05 | Treatment | -0.99 | -2.45 | 0.46 | fixed | | 0.74 | -1.33 | 0.18 |
| Green LED | m3 | Success ~ Days<br>* Treatment +<br>(1 \| Fish) | $\Delta AIC < 2$<br>and LRT p<br>> 0.05 | Days:<br>Treatment | -0.25 | -1.17 | 0.65 | fixed | | 0.46 | -0.55 | 0.57 |
| Green LED | m3 | Success ~ Days<br>* Treatment +<br>(1 \| Fish) | $\Delta AIC < 2$<br>and LRT p<br>> 0.05 | SD<br>(intercept) | 0.42 | | | random | Fish | | | |
| Warm LED | m3 | Success ~ Days<br>* Treatment +<br>(1 \| Fish) | $\Delta AIC < 2$<br>and LRT p<br>> 0.05 | (Intercept) | 2.66 | 1.84 | 3.48 | fixed | | 0.41 | 6.37 | 1.84E-10 |
| Warm LED | m3 | Success ~ Days<br>* Treatment +<br>(1 \| Fish) | $\Delta AIC < 2$<br>and LRT p<br>> 0.05 | Days | -0.17 | -0.75 | 0.40 | fixed | | 0.29 | -0.58 | 0.55 |
| Warm LED | m3 | Success ~ Days<br>* Treatment +<br>(1 \| Fish) | $\Delta AIC < 2$<br>and LRT p<br>> 0.05 | Treatment | -0.38 | -1.47 | 0.70 | fixed | | 0.55 | -0.68 | 0.49 |
| Warm<br>LED | m3 | Success ~ Days<br>* Treatment +<br>(1 \| Fish) | $\Delta$ AIC < 2<br>and LRT p<br>> 0.05 | Days:<br>Treatment | -0.56 | -1.29 | 0.17 | fixed | | 0.37 | -1.48 | 0.13 |
| Warm<br>LED | m3 | Success ~ Days<br>* Treatment +<br>(1 \| Fish) | $\Delta$ AIC < 2<br>and LRT p<br>> 0.05 | SD<br>(intercept) | 0.30 | | | random | Fish | | | |

## Notes

### Competing Interest Statement

The authors have declared no competing interest.

### Summary of Updates

We removed typos and added a junior author to the manuscript.

## References

Ali, M.A. (1975). Retinomotor responses. In Vision in fishes: new approaches in research, 313–355. Boston, MA: Springer US.

Appelbaum, L., Wang, G. X., Maro, G. S., Mori, R., Tovin, A., Marin, W., Yokogawa, T., Kawakami, K., Smith, S. J., Gothilf, Y., Mignot, E., & Mourrain, P. (2009). Sleep–wake regulation and hypocretin–melatonin interaction in zebrafish. Proceedings of the National Academy of Sciences, 106, 21942–21947. 10.1073/pnas.906637106

Ben-Moshe Livne, Z., Alon, S., Vallone, D., Bayleyen, Y., Tovin, A., Shainer, I., Nisembaum, L. G., Aviram, I., Smadja-Storz, S., Fuentes, M., Falcón, J., Eisenberg, E., Klein, D. C., Burgess, H. A., Foulkes, N. S., & Gothilf, Y. (2016). Genetically blocking the zebrafish pineal clock affects circadian behaviour. PLoS Genetics, 12, e1006445. 10.1371/journal.pgen.1006445

Ben-Moshe, Z., Alon, S., Mracek, P., Faigenbloom, L., Tovin, A., Vatine, G. D., Eisenberg, E., Foulkes, N. S., & Gothilf, Y. (2014). The light-induced transcriptome of the zebrafish pineal gland reveals complex regulation of the circadian clockwork by light. Nucleic Acids Research, 42, 3750–3767. 10.1093/nar/gkt1359

Boeck, P. D., Bakker, M., Zwitser, R., Nivard, M., Hofman, A., Tuerlinckx, F., & Partchev, I. (2011). The estimation of item response models with the lmer function from the lme4 package in R. Journal of Statistical Software, 39, 1–28.

Brüning, A., Hölker, F., Franke, S., Kleiner, W., & Kloas, W. (2018). Influence of light intensity and spectral composition of artificial light at night on melatonin rhythm and mRNA expression of gonadotropins in roach *Rutilus rutilus*. Fish Physiology and Biochemistry, 44, 1–12. 10.1007/s10695-017-0408-6

Burnside, B. (2001). Light and circadian regulation of retinomotor movement. In Progress in brain research (eds. Kolb, H., Ripps, H., and Wu, S.), 131, 477–485.

Cortesi, F., Tettamanti, V. & de Busserolles, F. (2022). The visual ecology of anemonefishes. In: Evolution, development and ecology of anemonefishes, 87–94.

Cortesi, F., Mitchell, L.J., Tettamanti, V., Fogg, L.G., de Busserolles, F., Cheney, K.L. & Marshall, N.J. (2020). Visual system diversity in coral reef fishes. Seminars in cell & developmental biology, 106, 31–42.

Cruz, L. M., Shillinger, G. L., Robinson, N. J., Tomillo, P. S., & Paladino, F. V. (2018). Effect of light intensity and wavelength on the in-water orientation of olive ridley turtle hatchlings. Journal of Experimental Marine Biology and Ecology, 505, 52–56. 10.1016/j.jembe.2018.05.002

Davies, T. W., Duffy, J. P., Bennie, J., & Gaston, K. J. (2016). Stemming the tide of light pollution encroaching into marine protected areas. Conservation Letters, 9, 164–171. 10.1111/conl.12191

De Russi, G., Montalbano, G., Gatto, E., Maggi, E., Cannicci, S., Bertolucci, C., & Lucon-Xiccato, T. (2024). Differential impact of artificial light at night on cognitive flexibility in visual and spatial reversal learning tasks. Animal Behaviour, 218, 173–183. 10.1016/j.anbehav.2024.10.008

Diamantopoulou, C., Christoforou, E., Dominoni, D. M., Kaiserli, E., Czyzewski, J., Mirzai, N., & Spatharis, S. (2021). Wavelength-dependent effects of artificial light at night on phytoplankton growth and community structure. Proceedings of the Royal Society B: Biological Sciences, 288, 20210525. 10.1098/rspb.2021.0525

Ferretti, M., Rossi, F., Benedetti-Cecchi, L., & Maggi, E. (2024). Ecological consequences of artificial light at night on coastal species in natural and artificial habitats: a review. Marine Biology, 172, 5. 10.1007/s00227-024-04568-2

Geoffroy, M., Langbehn, T., Priou, P., Varpe, Ø., Johnsen, G., Le Bris, A., Fisher, J. A. D., Daase, M., McKee, D., Cohen, J., & Berge, J. (2021). Pelagic organisms avoid white, blue, and red artificial light from scientific instruments. Scientific Reports, 11, 14941. 10.1038/s41598-021-94355-6

Jerlov, N. G. 1976. Marine Optics, 231. Amsterdam, the Netherlands; Oxford, UK; New York, NY: Elsevier Scientific Publishers. Elsevier Oceanography Series, 14.

Kieffer, J. D., & Colgan, P. W. (1992). The role of learning in fish behaviour. Reviews in Fish Biology and Fisheries, 2, 125–143. 10.1007/BF00042881

Lamb, T. D., Patel, H., Chuah, A., Natoli, R. C., Davies, W. I. L., Hart, N. S., Collin, S. P., & Hunt, D. M. (2016). Evolution of vertebrate phototransduction: cascade activation. Molecular Biology and Evolution, 33, 2064–2087. 10.1093/molbev/msw095

Laudet, V., & Ravasi, T. (Eds.). (2023). Evolution, development and ecology of anemonefishes: model organisms for marine science. Taylor & Francis. 10.1201/9781003125365

Li, L. (2019). Circadian vision in zebrafish: from molecule to cell and from neural network to behavior. Journal of Biological Rhythms, 34, 451–462. 10.1177/0748730419863917

Li, W., Zhang, D., Zou, Q., Bose, A. P. H., Jordan, A., McCallum, E. S., Bao, J., & Duan, M. (2024). Behavioural and transgenerational effects of artificial light at night (ALAN) of varying spectral compositions in zebrafish (*Danio rerio*). Science of The Total Environment, 954, 176336. 10.1016/j.scitotenv.2024.176336

Lucon-Xiccato, T., De Russi, G., Cannicci, S., Maggi, E., & Bertolucci, C. (2023). Embryonic exposure to artificial light at night impairs learning abilities and their covariance with behavioural traits in teleost fish. Biology Letters, 19, 20230436. 10.1098/rsbl.2023.0436

Marangoni, L. F. B., Davies, T., Smyth, T., Rodríguez, A., Hamann, M., Duarte, C., Pendoley, K., Berge, J., Maggi, E., & Levy, O. (2022). Impacts of artificial light at night in marine ecosystems—a review. Global Change Biology, 28, 5346–5367. 10.1111/gcb.16264

Marshall, N. J., Cortesi, F., de Busserolles, F., Siebeck, U. E., & Cheney, K. L. (2019). Colours and colour vision in reef fishes: past, present and future research directions. Journal of Fish Biology, 95, 5–38. 10.1111/jfb.13849

Mitchell, L. J., Cheney, K. L., Lührmann, M., Marshall, J., Michie, K., & Cortesi, F. (2021). Molecular evolution of ultraviolet visual opsins and spectral tuning of photoreceptors in anemonefishes (Amphiprioninae). Genome Biology and Evolution, 13, evab184. 10.1093/gbe/evab184

Mitchell, L.J., Phelan, A., Cortesi, F., Marshall, N.J., Chung, W.S., Osorio, D.C., & Cheney, K.L. (2024). Ultraviolet vision in anemonefish improves colour discrimination. Journal of Experimental Biology, 227, p.jeb247425.

Peña, M., Cabrera-Gámez, J., & Domínguez-Brito, A. C. (2020). Multi-frequency and light-avoiding characteristics of deep acoustic layers in the North Atlantic. Marine Environmental Research, 154, 104842. 10.1016/j.marenvres.2019.104842

Posit, T. (2025). RStudio: Integrated Development Environment for R. Posit Software [Computer software]. Posit. http://www.posit.co/

Powell, S. B., Mitchell, L. J., Phelan, A. M., Cortesi, F., Marshall, J., & Cheney, K. L. (2021). A five-channel LED display to investigate UV perception. Methods in Ecology and Evolution, 12, 602–607. 10.1111/2041-210X.13555

Sigurgeirsson, B., Þorsteinsson, H., Sigmundsdóttir, S., Lieder, R., Sveinsdóttir, H. S., Sigurjónsson, Ó. E., Halldórsson, B., & Karlsson, K. (2013). Sleep–wake dynamics under extended light and extended dark conditions in adult zebrafish. Behavioural Brain Research, 256, 377–390. 10.1016/j.bbr.2013.08.032

Smyth, T.J., Wright, A.E., McKee, D., Tidau, S., Tamir, R., Dubinsky, Z., Iluz, D. & Davies, T.W. (2021). A global atlas of artificial light at night under the sea. Elementa: Science of the Anthropocene, 9, 00049.

Sowersby, W., Kobayahsi, T., Awata, S., Sogawa, S., & Kohda, M. (2024). The influence of sleep disruption on learning and memory in fish. bioRxiv. 10.1101/2024.08.28.610197

Stanton, D. L., & Cowart, J. R. (2024). The effects of artificial light at night (ALAN) on the circadian biology of marine animals. Frontiers in Marine Science, 11, p.1372889. 10.3389/fmars.2024.1372889

Stieb, S.M., Cortesi, F., Jardim de Queiroz, L., Carleton, K.L., Seehausen, O., & Marshall, N.J. (2023). Long-wavelength-sensitive (*lws*) opsin gene expression, foraging and visual communication in coral reef fishes. Molecular Ecology, 32, 1656–1672.

Syposz, M., Padget, O., Willis, J., Van Doren, B. M., Gillies, N., Fayet, A. L., Wood, M. J., Alejo, A., & Guilford, T. (2021). Avoidance of different durations, colours and intensities of artificial light by adult seabirds. Scientific Reports, 11, p.18941. 10.1038/s41598-021-97986-x

Tidau, S., Smyth, T., McKee, D., Wiedenmann, J., D’Angelo, C., Wilcockson, D., Ellison, A., Grimmer, A. J., Jenkins, S. R., Widdicombe, S., Queirós, A. M., Talbot, E., Wright, A., & Davies, T. W. (2021). Marine artificial light at night: an empirical and technical guide. Methods in Ecology and Evolution, 12, 1588–1601. 10.1111/2041-210x.13653

Triki, Z., Fong, S., Amcoff, M., Vàsquez-Nilsson, S., & Kolm, N. (2023). Experimental expansion of relative telencephalon size improves the main executive function abilities in guppy. PNAS Nexus, 2, pgad129. 10.1093/pnasnexus/pgad129

Weschke, E. (2024). The nighttime ecology of coral reef fishes in a changing world. Doctoral Thesis, University of Bristol.

Zhang, M., Gao, X., Luo, Q., Lin, S., Lyu, M., Luo, X., Ke, C., & You, W. (2023). Ecological benefits of artificial light at night (ALAN): accelerating the development and metamorphosis of marine shellfish larvae. Science of The Total Environment, 903, 166683. 10.1016/j.scitotenv.2023.166683

Zhdanova, I. V., Wang, S. Y., Leclair, O. U., & Danilova, N. P. (2001). Melatonin promotes sleep-like state in zebrafish. Brain Research, 903, 263–268. 10.1016/S0006-8993(01)02444-1

